# Structural basis for Porcupine inhibition

**DOI:** 10.1101/2025.05.19.654776

**Authors:** Katrina A Black, Jesse I Mobbs, Hariprasad Venugopal, Toby A Dite, Andrew Leis, Lilian LL Wong, Laura F Dagley, David M Thal, Alisa Glukhova

**Affiliations:** The Walter and Eliza Hall Institute of Medical Research; Parkville, Australia; Department of Medical Biology, University of Melbourne, Melbourne, Victoria, Australia; Department of Cellular and Molecular Pharmacology, University of California, San Francisco, San Francisco, CA; Drug Discovery Biology, Monash Institute of Pharmaceutical Sciences, Monash University; Parkville, Australia; ARC Centre for Cryo-electron Microscopy of Membrane Proteins, Monash Institute of Pharmaceutical Sciences, Monash University; Parkville, Australia; Ramaciotti Centre for Cryo-Electron Microscopy, Monash University; Clayton, Australia; Department of Biochemistry and Pharmacology, The University of Melbourne; Melbourne, Australia

## Abstract

Wnt signalling is essential for embryonic development and tissue homeostasis, and its dysregulation is associated with multiple types of cancer. Porcupine (PORCN), an endoplasmic reticulum (ER)-resident membrane-bound O-acyltransferase, catalyses the palmitoylation of all 19 human Wnts—a critical modification required for their secretion and activity. This central role makes PORCN an attractive therapeutic target for Wnt-driven cancers, with several inhibitors currently in clinical trials. Here, we present high-resolution cryo-electron microscopy structures of human PORCN in complex with the inhibitors C59 (2.4 Å) and ETC159 (2.6 Å), as well as in a ligand-free state (3.3 Å). These structures reveal critical ordered water molecules that form a hydrogen-bonding network within the active site, mediating inhibitor binding. Our docking simulations of diverse PORCN inhibitors demonstrate that despite their different chemical scaffolds, these compounds adopt similar conformations within the acyl-CoA binding site and are also engaged through a conserved water molecule. Our findings provide a structural foundation for the rational design of next-generation PORCN inhibitors with improved pharmacological properties for cancer therapy.

## Introduction

Wnts are secreted glycoproteins that initiate a conserved signalling pathway critical for both embryonic development and adult tissue maintenance ^1^. Wnts undergo key post-translational modifications essential for their function, including the addition of a monounsaturated palmitoleate (16:1) to a conserved serine residue at the tip of Wnt “thumb” domain, a disulfide-bond stabilised β-hairpin ^2-4^. This acylation is catalysed by the enzyme Porcupine (PORCN), an ER-resident member of the membrane-bound O-acyltransferase (MBOAT) family^5^. Palmitoylation is essential for all 19 human Wnts, enabling their trafficking to the plasma membrane, secretion, and binding to Frizzled receptors ^2,3,6-8^. Through its universal role in Wnt processing, PORCN serves as a central gatekeeper for all Wnt-dependent signalling pathways ^6^.

Aberrant Wnt signalling drives the development of several cancers, including “Wnt-addicted” cancers that actively depend on the Wnt ligands for their growth. As PORCN represents a bottleneck in the secretion of all Wnt ligands and has no other established biological roles, it presents an attractive and potentially selective therapeutic target for Wnt-dependent cancers ^9-11^. Consequently, significant effort has been directed toward developing small-molecule inhibitors of PORCN ^12^. Several potent compounds have emerged from these efforts, with six reaching clinical trials: LGK974 (WNT974), ETC159, RXC004, RXC006 (AZD5055), XNW7201 and CGX1321 ^12,13^. These inhibitors share a common architecture of linearly connected hydrophobic rings, despite their diverse chemical scaffolds, and bind to PORCN with high affinity, often in the low nanomolar or even picomolar range ^13-16^.

The structural and mechanistic understanding of the entire MBOAT family and PORCN, in particular, has been limited until recently ^5^. The first structures of PORCN were reported in 2022 ^17^, providing insights into the overall molecular architecture of the enzyme, substrate binding, catalytic mechanism, and modes of inhibition. These structures were determined using cryo-electron microscopy (cryo-EM) with the aid of Fab fragments to facilitate particle alignment and image processing.

Here, we present a high-resolution structure of PORCN bound to the inhibitor C59 ^9,18^, determined at a resolution of 2.4 Å without the use of fiducial markers. The ordered component of this structure is 55 kDa, making it one of the smallest membrane proteins yet reported in this resolution range by cryo-EM. We have also determined structures of PORCN bound to ETC159 ^15^ at 2.6 Å and in a ligand-free state at 3.3 Å, providing a comprehensive structural characterisation of this important enzyme.

Our high-resolution structures provide critical insights into PORCN inhibitor binding, including an ordered water network that is essential for both inhibitor binding and catalysis. Combined with molecular docking studies, our analysis provides the determinants of inhibitor binding and selectivity. These findings establish a framework for structure-guided development of next-generation PORCN inhibitors with optimised pharmacological properties for cancer therapeutics.

## Results and discussion

### Cryo-EM of PORCN in the absence of fiducial markers

Despite the importance of MBOAT protein family members in lipid metabolism, their structural investigations are relatively recent, with the first X-ray structure of DltB published in 2018 ^19^. The challenges are their membrane localisation, their small size (<60 kDa), and the absence of large extra-membrane domains^5^. Many MBOAT family members form oligomers (e.g. SOAT1^20^, ACAT1^21,22^, DGAT1^23,24^) that facilitate their structure determination using cryo-EM. However, the Hedgehog morphogen acyltransferase, HHAT, the closest resolved homolog to PORCN at the commencement of this study, was monomeric and required the Fab or Nb binding to increase its size and serve as fiducial markers for particle alignment in cryo-EM ^25,26^.

Constant improvements in cryo-EM hardware and software continually push the boundaries of what is achievable ^27,28^. Thus, we reasoned that it should be possible to determine the PORCN structure in the absence of fiducial markers, even in its monomeric state. We chose to pursue isoform 3 (isoform C) of PORCN because it exhibited high acylation activity for most Wnts ^6^. To aid in expression, detection, and purification, our constructs contained a monomeric super-folded GFP and a StrepII tag at their C-terminus connected through a flexible linker ^29,30^. PORCN was expressed in Expi293 cells using the BacMam system ^31^ and purified using lauryl maltose neopentyl glycol (LMNG) detergent in the presence of POPS, which was previously shown to increase PORCN stability ^32^(**Figure S1A**). PORCN appeared to be monomeric and homogeneous based on multi-angle light scattering and negative stain imaging (**Figure S1B-C**).

The LMNG-solubilised PORCN was functional *in vitro*. We used liquid chromatography with tandem mass spectrometry (LC-MS) analysis to demonstrate the production of the palmitoylated peptide product, which is completely inhibited by the addition of the high-affinity PORCN inhibitor, C59^9^ (**Figure S1D-E**). We have also determined the K_m_ for PORCN-catalyzed palmitoylation of Wnt3a peptide using the recently developed fluorescence-based assay using 7-diethylamino-3-(4-maleimidophenyl)-4-methylcoumarin (CPM) dye^17^. The K_m_ for palmitoleoyl-CoA for LMNG-PORCN is 3 µm, which is slightly lower than the previously reported value (15 µM ^17^) (**Figure S1F**). These differences could be due to differences in PORCN isoforms (C vs A) or differences in purification conditions, such as the choice of detergent during purification or the absence of cholesterol hemisuccinate in our purification.

To identify the best small molecule for our structural studies, we employed CPM-based ThermoFluor assay^33^. In the absence of small molecules, the melting temperature (T_m_) of PORCN was 38.7 °C. The addition of PORCN inhibitors, C59^9^, GNF6231^34^, LGK974^14^ and ETC159^15^, increased the T_m_ of the PORCN to 43.9, 40.6, 40.6 and 40.0 °C, respectively. Notably, C59 stabilised PORCN by 5.2 °C and, thus, was initially chosen for structural studies (**Figure 1A**). The phospholipid 1-palmitoyl-2-oleoyl-sn-glycero-3-phospho-L-serine (POPS) was previously found to be essential for PORCN purification^32^. To confirm the requirement of POPS for PORCN stability, we also tested the effect of POPS on the T_m_ of PORCN. We purified PORCN in the absence of POPS and tested its thermostability with and without C59 and POPS. POPS increased the T_m_ of apo-PORCN by 2.4 °C and C59-bound PORCN by 5.2 °C (**Figure 1A**). Thus, POPS appears to elicit an effect on purified PORCN, suggesting it serves a stabilising role for detergent-purified protein.

**Figure 1.**
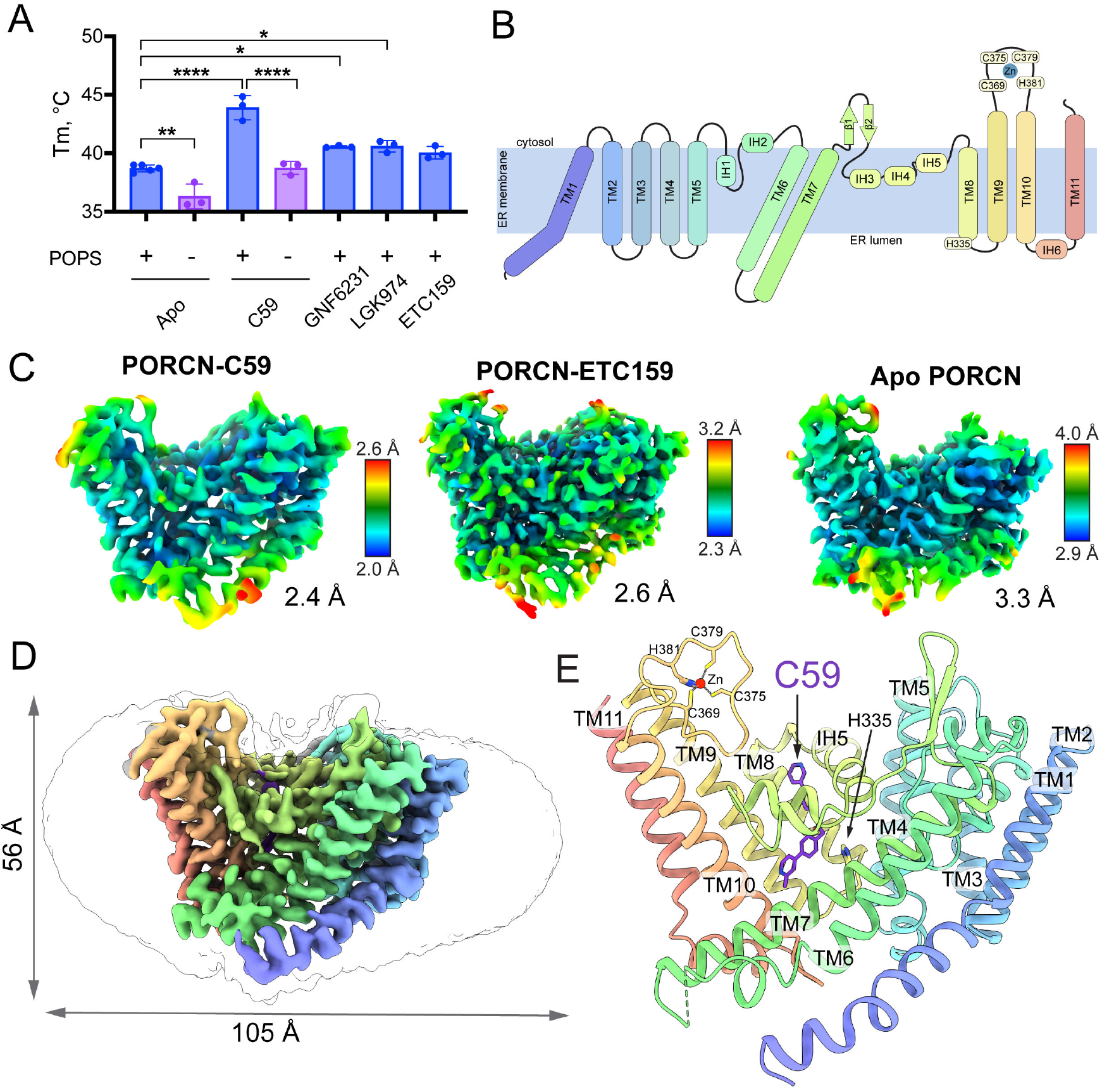
The overall architecture of PORCN. **A**) Thermal stability of PORCN in the presence of POPS and different inhibitors (n=3, mean ± SD, One-way ANOVA, *P < 0.05, **P < 0.01, ****P < 0.0001). **B)** PORCN topology diagram showing transmembrane helices (TM1-11, blue to red), intervening helices (IH1-6), and loops, catalytic histidine (H335) and the zinc-coordinating residues. **C)** Local resolution maps of PORCN-C59 (2.4 Å), PORCN-ETC159 (2.6 Å), and apo-PORCN (3.3 Å) determined by cryo-EM. **D)** Map of PORCN-C59 (map contour level 0.04) within the detergent micelle. **E)** Model of PORCN-C59 shown as a cartoon colored from N-terminus (blue) to C-terminus (red) with C59 inhibitor shown in purple sticks.

### A high-resolution cryo-EM structure of PORCN

We determined the structure of PORCN purified in LMNG in the presence of POPS and C59, ETC 159, and in a ligand-free state using cryo-EM to 2.4, 2.6, and 3.3 Å, respectively (**Figure 1B-E, S2-4**). The C-terminal GFP fusion did not contribute to particle alignment and was not visible during 2D classification (**Figure S2A**). Notably, we were able to resolve a 55 kDa membrane protein, almost entirely buried within a detergent micelle, to high resolution in the absence of any fiducial markers (**Figure 1C-D**).

While this manuscript was in preparation, Liu et al. published structures of PORCN with LGK974, Wnt3a peptide, and palmitoleoyl-CoA (resolutions 2.9-3.2 Å) determined by cryo-EM with the aid of a Fab^17^. Nevertheless, our PORCN-C59 represents the highest resolution inhibitor-bound PORCN structure determined to date and provides additional structural details that could aid future drug discovery efforts.

The structure of PORCN has a typical MBOAT fold and consists of 11 transmembrane helices, six short intervening-alpha helices and two beta strands (**Figure 1B, E**). Similar to other MBOAT family members, it resembles an hourglass, with a wider opening at the cytoplasmic site for Acyl-CoA binding and a smaller ER lumen-open site for binding Wnts. The conserved Zn^2+^ atom is coordinated by C369, C375, C379 and H381 (**Figure 1B, D**).

The C59-PORCN structure very closely resembles the PORCN-LGK974 structure^17^ (PDB 7URC) (RMSD 0.46 Å) (**Figure S5A-B**). The small deviations between the structures are in areas with high flexibility and poor map quality (e.g TM4-TM5 loop). Similar to the PORCN-LGK974 complex^17^, the ER-facing loops between TM6–TM7 (residues 222–229) and TM10–TM11 (residues 415–424) were unresolved. These loops lie in close proximity to Wnt in AlphaFold2-generated models of PORCN-Wnt3a complex and may only be resolved in full-length complexes of PORCN and Wnt (**Figure S6A**). Interestingly, the beginning of TM7 is the site of an alternative splicing in five PORCN isoforms expressed in humans^35^(**Figure S6B**). This loop faces the poorly conserved L1 Wnt loop ^36^, and thus may be responsible for PORCN-isoform-mediated selectivity^6^ (**Figure S6C**). In addition, the shorter TM7 and the reduced buried PORCN-Wnt contact area may be responsible for the lower activity of PORCN isoform A, as observed previously^6^.

For the first time, the high resolution of our C59-PORCN structure allowed the observation of several ordered water molecules. These included several structural waters that bridge neighbouring secondary structure elements through H-bonds (**Figure S2C**) and water molecules important for ligand binding (**Figure 2A-C** and see below). Importantly, the structural water near residue N300 likely stabilises the PORCN active site by bridging and stabilising the IH5 and IH1-IH2 loop (**Figure S2C-D**). This interaction is likely responsible for the significant loss of activity in the equivalent N301A mutant^32^.

**Figure 2.**
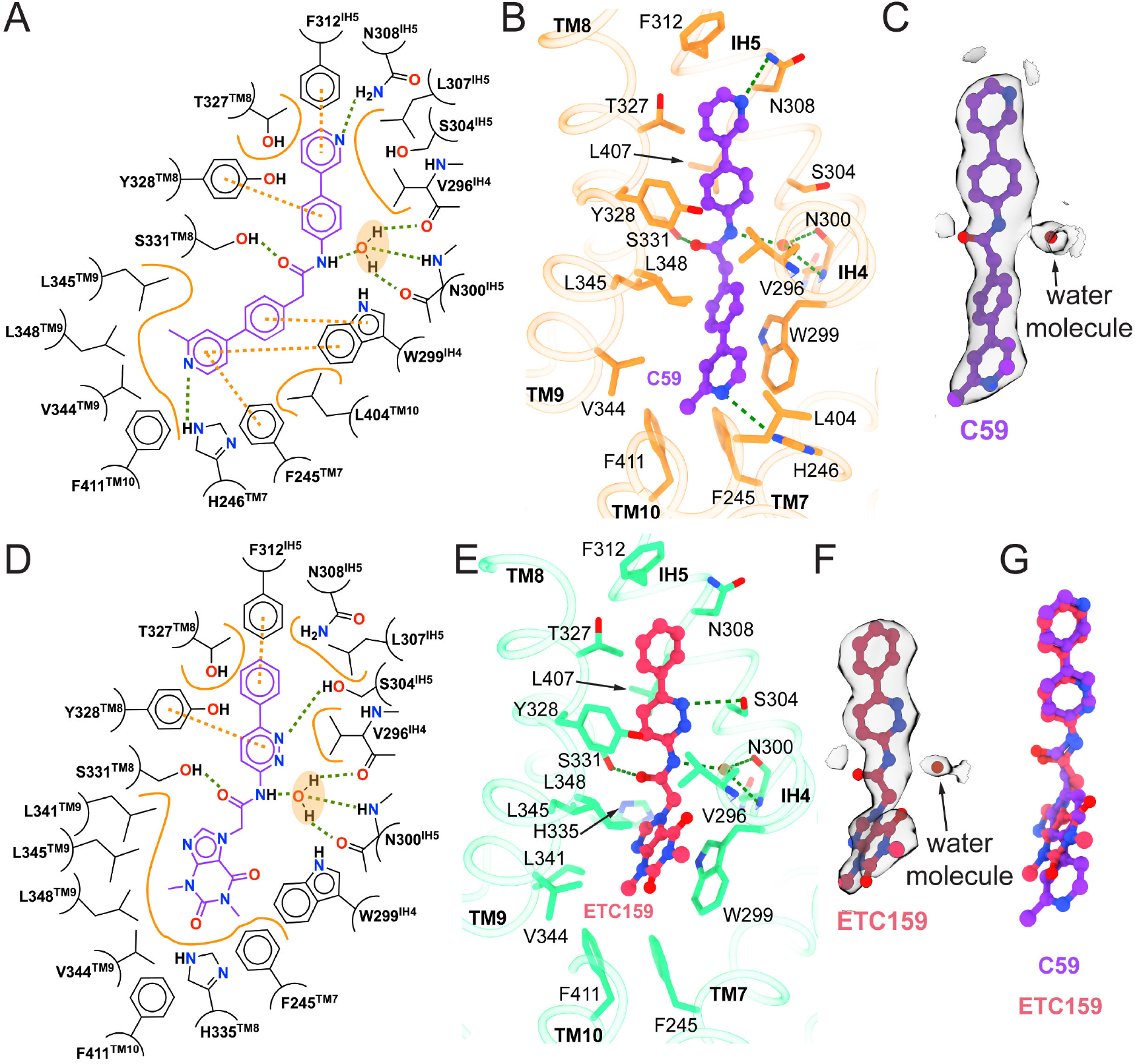
C59 and ETC inhibitor binding to PORCN. **A)** Schematic representation of interactions between PORCN and C59 (purple). **B)** Cartoon representation of C59 binding site. **C)** Cryo-EM density map showing C59 and an ordered water molecule (map contour level 0.034). **D)** Schematic representation of interactions between PORCN and ETC159 (pink). **E)** Cartoon representation of ETC159 binding site. **F)** Cryo-EM density map showing ETC159 and an ordered water molecule (map contour level 0.034). **G)** Superposition of C59 and ETC159 in the PORCN binding site. H-bonds are shown as green, π-stacking as orange dashed lines, and Van der Waals interactions as solid orange lines. Key interacting residues are shown as sticks, ligands in ball-and-stick representation, water as a red sphere, and the cryo-EM map is displayed as a transparent grey surface.

### The binding mode of the potent PORCN inhibitors C59 and ETC159

C59 and ETC159 bind in the same pocket as a closely related LGK974 in the PORCN-LGK974 structure (PDB 7URC), adopting an L-shape within the cytoplasmic acyl-CoA-binding site ^17^ defined by helices IH4, IH5 and TM7, TM8 and TM9 and next to a catalytic H335 (**Figure 1E** and **2A-G**). Key interactions involved in inhibitor binding include an H-bond with S331, van der Waals interactions between the aromatic groups of the inhibitors and the hydrophobic side chains lining the pocket, as well as π-π and π-edge interactions with W299, F245, F312, and Y328. In PORCN-C59, the binding was further stabilised by H-bonds with N308 and H246 (**Figure 2A-B**). In PORCN-ETC159, an additional H-bond was formed with S304 (**Figure 2D-E**).

An ordered water molecule stabilises C59 and ETC159 by acting as a bridge between the inhibitors’ amide nitrogen and the amide and carbonyl groups of N300 and the carbonyl of V296 (**Figure 2C and F**). Since most PORCN inhibitors contain an amide group separating heterocyclic rings ^13,37^, this water likely forms crucial H-bond interactions, contributing significantly to the inhibitor’s binding energy.

While not resolved due to lower resolution in acyl-CoA-bound structure in Liu *et al*. (PDB 7URA) ^17^, this water likely stabilised the palmitoleoyl-CoA in the “curled up” conformation, bringing the carbonyl oxygen of the fatty acid in the vicinity of catalytic H335 for the acylation reaction (**Figure S5C**). Thus, this water molecule is crucial for positioning PORCN substrates in a conformation suitable for acyl group transfer.

We also observed a lipid-like moiety capping the PORCN binding site at the cytosolic site with the acyl tails projecting towards the active site (**Figure S7A**). This density overlaps with the phospho-adenosine headgroup of the palmitoleoyl-CoA (in PDB 7URA) and caps the PORCN binding site, likely making van der Waals interactions with the inhibitor (**Figure S7B**). Unfortunately, we were unable to unambiguously identify this molecule. However, it is tempting to speculate that this might be a binding site for a lipid, that co-purified with PORCN or POPS that was added during purification to aid with PORCN stability^32^ (**Figure 1A**). Interestingly, this density is present in all inhibitor-bound PORCN structures determined to date (this manuscript and in Liu *et al*.^17^).

### PORCN conformational change upon inhibitor binding

To understand the conformational change upon inhibitor binding, we compared the PORCN structure determined in the absence of inhibitors with that of PORCN-C59 and PORCN-ETC159 (**Figure 3**). All three structures were nearly identical with root mean square deviations (RMSDs) <0.5 Å. There are minimal differences in TM positions or the loops, including those that define the PORCN binding site (**Figure 3A**). Contrary to conformational changes observed in HHAT upon IMP-1575 binding^26^, the conformation of the amino acid residues lining the binding site are almost identical, with the exception of a small rotation of W299 in the ETC159-bound structure, which aligns parallel to the di-oxo-purine group of ETC159 for π-π stacking interactions (**Figure 3B**). Such surprising invariance between apo and inhibitor states is likely due to the nature of the binding pocket that presents a long hydrophobic channel across the ER membrane for transferring palmitoleoyl-CoA from the cytoplasm into the lumen of the ER for attachment to Wnt.

**Figure 3.**
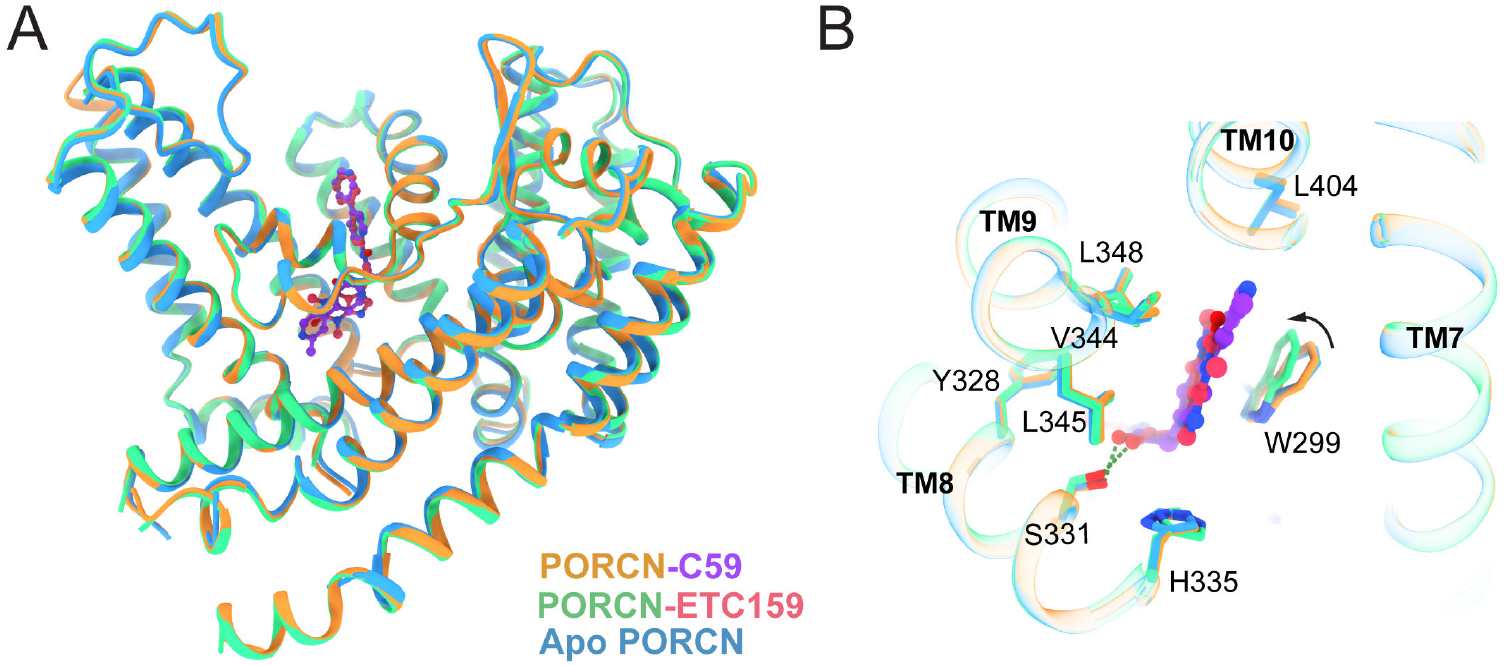
Comparison between apo and inhibitor-bound PORCN structures. **A)** Superposition of PORCN-C59 (orange-purple), PORCN-ETC159 (green-pink), and apo PORCN (blue) structures. **B)** Close-up view of the inhibitor binding pocket showing minimal conformational changes, with the exception of W299 in the ETC159-bound structure, which rotates to form π-π stacking interactions with the inhibitor. H-bonds are shown as green dashed lines.

However, an alternative explanation for the apparent lack of conformational change upon inhibitor binding might be the partial occupancy of the PORCN binding site with endogenous lipids or acyl-CoA molecules. The apo-structure exhibits weak electron density in the binding pocket (**Figure S8**), less pronounced than in our inhibitor-bound structures or the acyl-CoA-bound structure reported by Liu *et al*. ^17^ (PDB 7URA). We interpreted this as water molecules within the acyl-CoA transfer channel, based on PORCN molecular dynamics simulations ^38^. Alternatively, this density might instead represent a low-occupancy endogenous ligand present during purification.

### Docking of PORCN inhibitors with high affinity

We used our high-resolution PORCN structure to conduct docking analyses to understand the binding modes of PORCN inhibitors with diverse chemical scaffolds.

First, we tested the performance of our docking simulations using C59, ETC159, and LGK974 to compare how simulations recapitulate experimentally validated poses^17^ (**Figure S9A-F**). We performed docking both in the absence and presence of water molecules to evaluate their contribution to binding. All three ligands adopted similar poses to their respective structures. However, docking in the presence of water molecules yielded a better agreement with experimental data, due to the anchoring of the acetamide group through hydrogen bonds to S331 and N300, facilitated by a bridging water molecule. Thus, we used the PORCN-C59 structure with modelled water molecules, but with the ligand removed, to dock several compounds selected for their clinical relevance (RXC004^16^) and the overall diversity of their chemical scaffolds (GNF6231 ^34^, IWP-O1 ^39^, IWP-L6 ^40^) (**Figure 4**).

**Figure 4.**
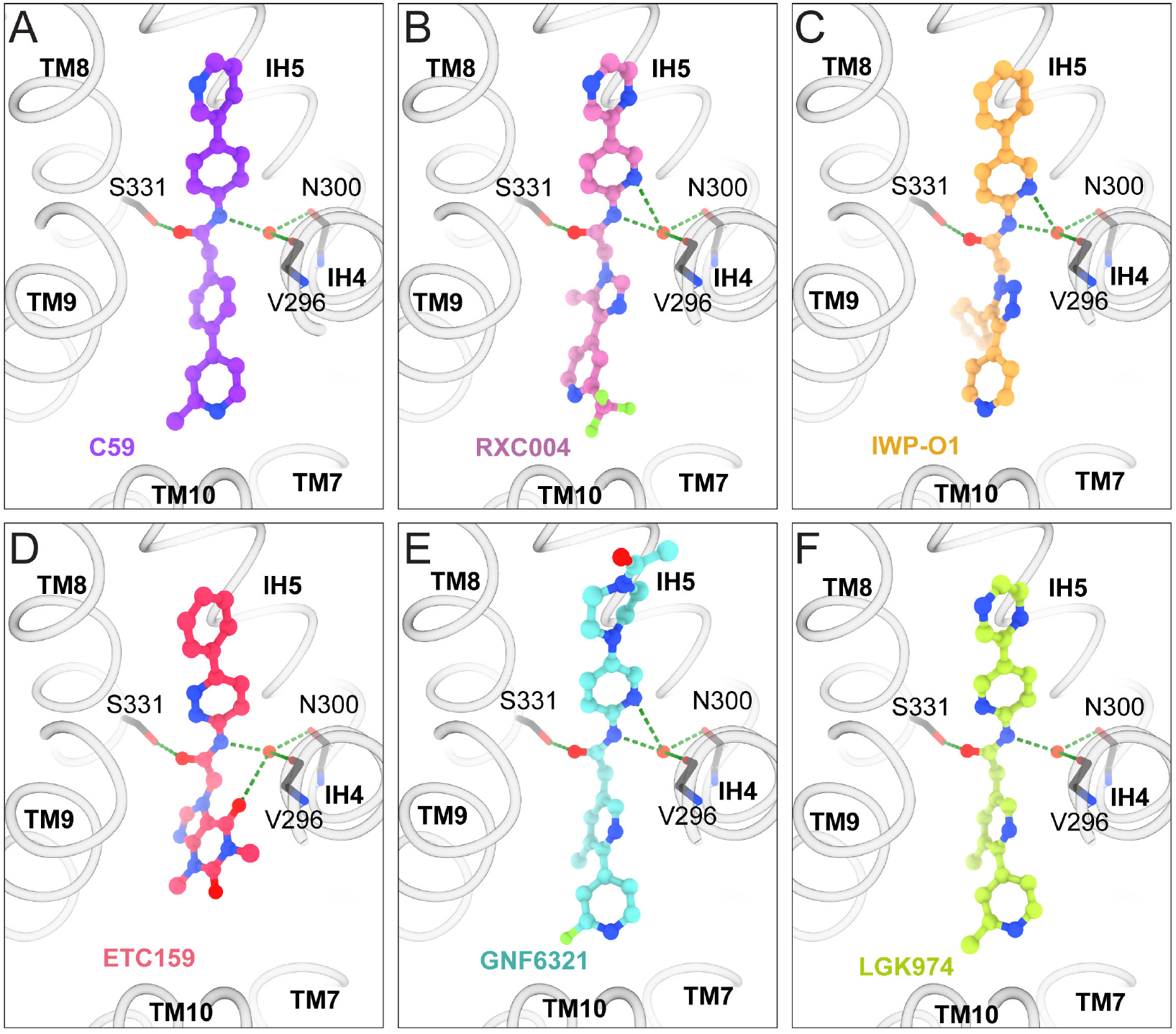
Molecular docking of PORCN inhibitors. **A-F)** Docking poses of diverse PORCN inhibitors: C59 (**A**, purple), RXC004 (**B**, pink), IWP-O1 (**C**, orange), ETC159 (**D**, red), GNF6231 (**E**, cyan), and LGK974 (**F**, light green). All inhibitors adopt a similar L-shaped conformation in the binding site, anchored by hydrogen bonds (green dashed lines) with S331 and N300, mediated by a water molecule.

All compounds bound in the narrow acyl-CoA binding site and generally overlapped with C59, adopting an L-shape to conform to the chemical pocket while making expected interactions with their acetamide groups through hydrogen bonds to S331 and N300 that was mediated by a water molecule. The two linear aromatic rings occupy positions distal to the Wnt binding site and were constrained by the tight hydrophobic channel formed by IH4, IH5, TM8, and TM9. Most variability in compound binding poses occurs at the luminal side of PORCN, where the binding site widens to accommodate the curling of acyl-CoA upon its attachment to Wnt. This area is spacious enough to accommodate linear cyclic arrangements of C59, LGK974, and GNF6231, as well as the dioxopurine group of ETC159, the trifluoromethylpyridine group of RXC004, and the branched phenyl-pyridine-triazole group of IWP-O1 (**Figure 4**).

Unexpectedly, the initial docking simulations revealed a unique pose for IWP-L6 at the entrance of the binding site (**Figure S9G**). However, this outlier was likely due to steric clashes between the bulky IWP-L6 structure and side chains within the binding site. Upon relaxing the side chain conformations through induced fit docking, IWP-L6 adopted the expected binding pose (**Figure S9H**), revealing the anticipated network of hydrogen bonds and the phenyl-tetrahydrothienopyrimidine group positioned within the wider region of the binding site.

Overall, despite the diverse chemical scaffolds of PORCN inhibitors, these compounds adopt similar binding poses and appear to compete for the acyl-CoA binding site but likely do not prevent Wnt binding, in agreement with the structure of Wnt3a-peptide simultaneously bound with LGK974 (PDB 7URD) ^17^. The structure, together with our docking simulations, provides explanations for the similarities in chemical scaffolds of effective PORCN inhibitors.

## Conclusions

Our high-resolution cryo-EM structures of PORCN provide insights into the binding mechanism of potent PORCN inhibitors. Notably, we achieved a 2.4 Å resolution structure of this 55 kDa membrane protein without the aid of fiducial markers, demonstrating the advancing capabilities of cryo-EM for small membrane proteins. As with similarly reported structures (e.g., GAT1), relatively large initial particle stacks and high on-grid concentrations appear to correlate with improved likelihood of structure determination for this class of proteins. Our high-resolution structures revealed a critical ordered water molecule within the active site that mediates key interactions with PORCN inhibitors.

The structural analysis of C59 and ETC159 binding, complemented by our docking studies of diverse PORCN inhibitors, reveals the common binding architecture required for high-affinity inhibition. Despite their chemical diversity, these compounds adopt similar L-shaped conformations within the acyl-CoA binding site and engage in conserved interactions, particularly through their acetamide groups. The wider pocket at the luminal side accommodates various chemical moieties, explaining the structural diversity observed among effective PORCN inhibitors.

The remarkable conformational similarity between apo and inhibitor-bound structures suggests that PORCN maintains a relatively rigid hydrophobic binding channel optimised for transferring palmitoleoyl-CoA across the ER membrane.

The structural insights from this work advance our understanding of PORCN’s role in Wnt signalling and provide valuable insights for targeting Wnt signalling in cancer therapeutics. As five PORCN inhibitors have already progressed to clinical trials, these structural details may facilitate the development of compounds with improved potency, selectivity, and physicochemical properties.

## Methods

### Cell culture

*Spodoptera frugiperda* Sf9 cells were cultured in ES-900 media (Expression Systems) at 27°C. Expi293 Freestyle cells were cultured in Expi293 Freestyle medium at 37°C.

### Protein expression and purification

cDNA encoding isoform 3 (isoform C) of human PORCN (with C187A mutation to curb S-palmitoylation^41^) was cloned into a pEG BacMam vector and fused to a C-terminal green fluorescent protein (GFP ^30^) followed by a StrepII tag and separated by a human rhinovirus (HRV)-3C protease cleavage site. Heterologous expression of PORCN was carried out using the BacMam system, as previously described ^31^. Bacmids were generated by transforming constructs into DH10Bac cells before baculoviruses were produced by transfecting Sf9 cells with the bacmid using Fugene HT (Promega). After two rounds of amplification, baculoviruses were used to transduce Expi293 cells for expression. Suspension cultures were grown at 37°C to a density of approximately 2.5 million cells/mL and then infected with 10% (v/v) baculovirus. Cells were harvested exactly 48 hours post-transduction, flash-frozen in liquid N_2_ and stored at -80°C for future purification.

The pellet from 1.2 L of cell culture was resuspended in 240 mL lysis buffer (40 mM HEPES pH 8.0, 350 mM NaCl, 30% glycerol (w/v), 10 mM MgCl_2_, benzonase, Roche cOmplete protease inhibitor tablets, 1.0% LMNG (w/v) and 100 uL BioLock (IBA). The cell pellet was lysed by dounce homogenisation with ice-cold pestle B before extraction at 4°C for 2 hours whilst nutating. The lysate was clarified by centrifugation for 30 minutes at 21,000 x g at 4°C before filtration of the supernatant through a glass fibre filter under vacuum. The supernatant was applied to 2.5 mL StrepTactin XT resin (IBA) that was equilibrated in W1 buffer (30 mM HEPES pH 7.5, 350 mM NaCl, 15% (w/v) glycerol, 0.3% LMNG) and loaded into a glass column. The supernatant was incubated with resin for 1 hr at 4°C. The beads were washed with a series of gentle exchange buffers (15 mL each) generated from varying ratios (50:50, 25:75, 10:90, 5:95, 1:99) of W1 buffer and W2 buffer (20 mM HEPES pH 8.0, 350 mM NaCl, 0.01% LMNG). PORCN was eluted in 6 equivalents to the bed volume of elution buffer (20 mM HEPES pH 8.0, 350 mM NaCl, 0.01% LMNG, 0.1 mgml^-1^, 50 mM biotin). The eluent was concentrated to 500 µL and purified by SEC (Superdex 200 10/300 GL column (Cytiva)) in buffer with 20 mM HEPES, 150 mM NaCl, 0.01% LMNG, pH 8.0). For the purification of PORCN in the presence of inhibitors, C59 or ETC159 was added at a concentration of 150 nM to all buffers.

### Negative stain EM imaging

Samples were diluted to 0.008 mg/ml and applied to a glow-discharged carbon-coated Cu grid (200 mesh). After 60 s the solution was blotted off and the grid was stained in 0.8% uranyl formate solution for 30 s, followed by blotting off the excess solution and two washes with water. The grids were imaged at room temperature using a Talos L120C transmission electron microscope (Thermo Fisher Scientific, USA) at a nominal magnification of 52,000x corresponding to a pixel size of 2.4 Å, and a defocus value of ∼-1 *µ*m.

### Grid preparation and cryo-EM imaging

PORCN samples (35 mg/ml) copurified in the absence of inhibitors or presence of C59 or ETC159 were applied to UltrAufoil 1.2/1.3 300 mesh holey grids (Quantifoil GmbH, Großlöbichau, Germany), that were freshly glow discharged in air at 15 mA for 180 s using a Pelco EasyGlow. 3 µl of sample was applied to the grids at 4°C and 100% humidity and plunge-frozen in liquid ethane using a Vitrobot Mark IV (Thermo Fisher Scientific, USA) with a blot time of 2s and blot force of 10. The Titan Krios was operated at an accelerating voltage of 300 kV with a 50 μm C2 aperture, 100 µm objective aperture inserted, and zero-loss filtering with a slit width of 10 eV, at an indicated magnification (**Table S1**) in nanoprobe EFTEM mode. Data were collected using aberration-free image shift (AFIS) with Thermo Fisher EPU software. Additional parameters for each dataset are listed in **Table S1**.

### Cryo-EM image processing and model building

#### PORCN-C59 dataset

10,069 movies at 0.65 Å/pix were motion corrected using Patch Motion Correction in cryoSPARC v3.3.2 ^42^, followed by Patch CTF Estimation (**Figure S2**). Initially, particles were picked using Blob picker, followed by 2D classification, heterogeneous and non-uniform refinement^43^ in cryoSPARC. This map was used to generate templates for template-based picking in the next round of data processing. A total of 11.8 million particles were picked, extracted (4× binned, 80 pix box, 2.6 Å/pix) and subjected to 3 rounds of heterogeneous refinement against the previously generated PORCN map and a “junk” class. The resulting 3.7 million particles were re-extracted to improve their centering (4x binned, 80 pix box, 2.6 Å/pix) and subjected to one round of 2D classification, followed by re-extraction (2x binned, 192 pix box, 1.3 Å/pix) and another two rounds of 2D classification and two rounds of heterogeneous refinement. The resulting 1,048,916 particles were extracted at full resolution (384 pix box, 0.65 Å/pix) and subjected to Non-Uniform Refinement ^43^, Reference-Based Motion Correction followed by another round of Non-Uniform Refinement. We generated a mask around the protein using EMAN2^44^ and used it for local refinement, yielding a final map at 2.4 Å (FSC = 0.143).

#### PORCN-ETC159 dataset

10,714 movies at 0.65 Å/pix were collected and processed similarly to the PORCN-C59 dataset in cryoSPARC v4.5.1 ^42^ with minor modifications (**Figure S3**). The final dataset contained 92,229 particles that yielded a map at 2.6 Å resolution (FSC = 0.143).

#### Apo PORCN dataset

10,714 movies at 0.65 Å/pix were collected and processed similarly to the PORCN-C59 dataset in cryoSPARC v4.5.1 ^42^ with minor modifications (**Figure S3**). The final dataset contained 92,229 particles that yielded a map at 3.3 Å resolution (FSC = 0.143).

#### Modelling

For modelling, the PORCN isoform C model generated using AlphaFold2^45^ (for PORCN-C59 modelling) or the PORCN-C59 model (for PORCN-ETC159 and Apo-PORCN structures) was rigid-body fitted into the density maps using UCSF ChimeraX^46^ and subjected to repeated rounds of manual model building in COOT^47^ and real-space refinement in PHENIX^48^. The C59 and ETC159 inhibitors were identified and manually built into the relevant maps. Both inhibitors were generated from SMILES codes, and geometry restraints were generated using the GRADE webserver^49^. Lastly, the models were quality assessed using MolProbity^50^ before PDB deposition. The figures were generated using UCSF ChimeraX^46^.

### SEC-MALS

Size-exclusion chromatography coupled to multi-angle light scattering (SEC-MALS) was performed using a X-bridge Protein BEH SEC column, 3.5 µm, 7.8 mm x 150 mm (Waters) coupled with a DAWN HELEOS II light scattering detector (Wyatt Technology, USA) and an Optilab T-rEX refractive index detector (Wyatt Technology, USA). The system was equilibrated in 20 mM HEPES (pH 8.0), 150 mM NaCl, 0.01% LMNG and calibrated with BSA (2 mg/mL). 20 µL of purified PORCN at 1.3 mgml^-1^. All data were collected and processed using ASTRA software (v7.3.1.9, Wyatt Technology).

### CPM Thermal Shift Assay

A CPM thermal shift assay was performed to evaluate PORCN protein stability under various conditions, including PORCN alone, PORCN without POPS, and PORCN in the presence of various inhibitors (C59, LGK-974, GNF6231 and ETC159). PORCN without POPS was carried out by purification of PORCN in the absence of exogenous POPS added during the purification; in all other conditions, POPS was added during PORCN purification, but no additional POPS was added during the assay. PORCN was assayed at a final concentration of 0.01 mg/ml and each inhibitor was present at a final concentration of 1 μM. Each condition was measured in triplicate. The assay was carried out in 30 mM HEPES pH 7.0, 100 mM NaCl and 0.01% LMNG. For each sample, 5 μL of 0.04 mg/mL (4×) protein was combined with 5 μL of 4× drug solution (inhibitors dissolved in the assay buffer) in a microplate well, followed by centrifugation at 1000×g for 1 minute. Then, 10 μL of 20 μM (2×) CPM stock solution was added to achieve a final concentration of 10 μM in a 20 μL reaction volume, followed by another centrifugation step at 1000×g for 1 minute. To prevent evaporation, 4 μL of silicone oil was layered on top of each sample, followed by centrifugation at 1000×g for 1 minute. All samples were incubated at room temperature for 30 minutes before fluorescence measurements were taken with excitation at 380 nm and emission at 470 nm during a temperature ramp from 5°C to 85°C at a rate of 1°C/minute. Each experimental condition had a matched control without protein (i.e., CPM alone, or CPM in the presence of drug) for subtraction of baseline CPM signal. Raw data were imported into GraphPad Prism (v 9.5.0), each sample was baseline corrected with its matching CPM control, and a Boltzmann sigmoidal curve was fitted to the corrected data. The midpoint of the transition represents the protein melting temperature (T_m_).

### LC-MS activity assay

The palmitoleoylation of WNT3A peptide was assessed using an intact peptide LC/MS assay. The reaction mixture (20 μL total volume) contained PORCN (7.5 μM final concentration), WNT3Ap peptide ((50 μM final concentration, sequence MHLKCKCHGLSGSCEVKTCWW with disulfide bonds between C5-C19 and C7-C14), freshly prepared palmitoleoyl CoA (100 μM in 20 mM HEPES pH 8.0), and reaction buffer (20 mM HEPES pH 8.0, 100 mM NaCl, 0.01% LMNG, 0.1 mg/mL POPS). The reaction was incubated at 15°C for 2 hours in a PCR thermal cycler.

Samples were cleaned up using C18 STAGE tips and collected into new tubes by centrifugation. The collected peptides were lyophilized to dryness using a CentriVap (Labconco), before reconstituting in 10 µl 0.1% FA/2% ACN ready for mass spectrometry analysis.

Peptides were separated by reverse-phase chromatography on a 25cm ProSwift RP-4H monolith column (Thermo Scientific) using a micro-flow HPLC (Ultimate 3000). The HPLC was coupled to a Maxis II Q-TOF mass spectrometer (Bruker) equipped with an ESI source. Peptides were loaded directly onto the column for online buffer exchange at a constant flow rate of 50 µl/min with buffer A (99.9% Milli-Q water, 0.1% FA) using a valve to direct flow to waste. After 10 minutes, the valve was switched to direct flow to the mass spectrometer, and peptides were eluted with a 10-min linear gradient from 2 to 90% buffer B (99.9% ACN, 0.1% FA). The Maxis II Q-TOF was operated in full MS mode using Compass Hystar 5.1. Settings for the 40 samples per day method were as follows: Mass Range 100 to 3000m/z, Capillary Voltage 4500V, Dry Gas 4 l/min, Dry Temp 200°C. Data was analysed with DataAnalysis version 5.2 (Bruker) and peptide masses were deconvoluted using the maximum entropy method. The masses of the unmodified and palmitoylated peptides are 2431.9 and 2668.1, respectively.

### CPM activity assay - Katrina

The activity of purified PORCN was assessed using a CPM-based fluorescence assay. When PORCN transfers palmitoleate to Wnt, the liberated CoA’s sulfhydryl group interacts with CPM to produce fluorescence. Experiments were performed in 96-well black-bottom plates (Greiner) with each condition tested in triplicate. The reaction mixture containing CPM solution (50 μM in 20 mM HEPES pH 8.0, 100 mM NaCl, 0.01% LMNG), WNT3Ap peptide (50 μM final concentration, sequence MHLKCKCHGLSGSCEVKTCWW with disulfide bonds between C5-C19 and C7-C14, manufactured by ChinaPeptides), freshly prepared palmitoleoyl-CoA (at specific concentrations of 2.5, 5, 10, 30, and 60 μM), and 0.1 mg/ml POPS was first prepared and added to the plate. After incubation at room temperature, reactions were initiated by adding PORCN (1 μM final concentration) and proceeded for 30 minutes at 37°C. CPM fluorescence was measured with an excitation of 355 nm and emission of 460 nm. Michaelis-Menten kinetic analysis was performed using GraphPad Prism 9.

### Computational molecular docking

The PORCN-C59 complex model was used for docking. The C59 ligand was removed, and the protein was prepared using the Schrödinger Protein Preparation Wizard. This process adds hydrogen atoms, assigns bond orders, optimises the hydrogen-bonding network, and performs a restrained minimisation using the OPLS4 force field. Grid generation was performed based on the location of C59 with default settings. These processes were performed with and without the central water in the ligand binding site.

The ligands were prepared from SMILES using Schrödinger LigPrep, to generate 3D structures, different ionisation states and tautomeric forms at a physiological pH of 7.4. The ligand structures were minimised using the OPLS4 force field. The ligands were docked using the default settings of Schrödinger Glide extra precision docking (XP).

The induced fit docking protocol of Schrödinger was used to further model the binding pose of IWP-L6. This protocol uses a combination of Glide XP and Prime to accurately model ligand and protein side chains. Preparation was performed similarly to previously, a grid box was built with the centroid of the C59 ligand, Prime refinement of side chains within 5 Å, and a maximum of 20 poses were redocked with Glide XP.

## Figure legends

**Supplementary Figure 1. Purification and functional characterisation of PORCN. A)** Size exclusion chromatography profile of purified PORCN with SDS-PAGE analysis of peak fractions (inset). **B)** SEC-MALS analysis of PORCN showing a monodisperse peak at 170 kDa. **C)** Representative negative stain EM images of purified PORCN-GFP. **D)** LC-MS analysis of PORCN-catalysed palmitoylation of Wnt3a peptide in the absence (middle) or presence (bottom) of C59 inhibitor, compared to no PORCN control (top). **E)** Quantification of LC-MS data (n=3, mean ± SEM, One-way ANOVA, *P < 0.05, **P < 0.01, n.s. not significant). **F)** Kinetics of PORCN-dependent Wnt3a peptide acylation. Data fitted to the Michaelis-Menten equation (K_m_ = 3 μM).

**Supplementary Figure 2. Cryo-EM data processing of PORCN-C59 complex. A)** Data processing workflow for PORCN-C59 complex. **B)** Gold-standard FSC curves for the final 3D reconstruction with different masking strategies. **C-D)** Cryo-EM density (transparent grey surface, map contour level 0.02) for the structural water molecules (red spheres) in PORCN-C59 structure. Coordinating residues are shown as sticks. H-bonds are shown as green dashed lines.

**Supplementary Figure 3. Cryo-EM data processing of PORCN-ETC159 complex. A)** Data processing workflow for PORCN-ETC159 complex. **B)** Gold-standard FSC curves for the final 3D reconstruction with different masking strategies.

**Supplementary Figure 4. Cryo-EM data processing of apo-PORCN. A)** Data processing workflow for apo-PORCN. **B)** Gold-standard FSC curves for the final 3D reconstruction with different masking strategies.

**Supplementary Figure 5. Structural comparison with published PORCN structures. A)** Superposition of PORCN-C59 (orange-purple) and PORCN-LGK974-Fab (PDB 7URC, blue-light green-grey) structures. **B)** Close-up view of superimposed C59 and LGK974. **C)** Overlay of the PORCN-acyl-CoA (PDB 7URA, blue-light green) with the PORCN-C59 structures (C59 omitted for clarity) showing that the water molecule (red sphere) observed in our PORCN-inhibitor structures is important for positioning the acyl chain for catalysis.

**Supplementary Figure 6. Models of 5 PORCN isoforms and their predicted interactions with Wnts. A)** AlphaFold3-generated models of five PORCN isoforms in complex with Wnt3a superposed with the PORCN-C59 structure. The ER lumen-facing loops (TM6-TM7 and TM10-TM11) that are unresolved in our structures are predicted to interact with Wnt. **B)** Sequence alignment of five human PORCN isoforms showing the alternatively spliced region at TM7. **C)** Model of PORCN-Wnt3a complex coloured by sequence conservation among human Wnts, highlighting the variable L1 loop region in Wnt that may contribute to PORCN isoform selectivity.

**Supplementary Figure 7. Lipid-like density at the entrance to the PORCN inhibitor binding site. A)** Unassigned lipid-like density (transparent grey surface, map contour level 0.048) observed at the cytosolic entrance of the PORCN-C59 binding pocket. **B)** Superposition with acyl-CoA from PDB 7URA (light green) showing the lipid-like density overlapping with the phospho-adenosine headgroup of acyl-CoA.

**Supplementary Figure 8. Electron density in the binding site of apo-PORCN**. Weak electron density (transparent grey surface, map contour level 0.034) observed in the binding site of apo-PORCN, which could represent water molecules or low-occupancy endogenous ligands.

**Supplementary Figure 9. Docking validation and analysis of IWP-L6 binding. A-F)**, Comparison of experimentally determined (purple, pink or light green) and docked (grey or dark grey) poses of C59, ETC159, and LGK974, showing improved accuracy when the water molecule is included in the docking simulations. **G)** Initial rigid protein docking of IWP-L6 (orange) showing binding at the entrance of the binding site. **H)** Induced fit docking of IWP-L6 showing the expected binding mode after allowing protein side-chain flexibility.

## Acknowledgements

This work was funded through a CSL Centenary Fellowship (AG) and the estate of Akos and Marjorie Talon (AG). DMT was funded by a National Health and Medical Research Council of Australia Investigator Grant (APP1196951). We acknowledge the use of the facilities at the Monash University Ramaciotti Centre for cryo-electron microscopy, the Ian Holmes Imaging Centre, Bio21 Institute, the Monash University MASSIVE high-performance computing facility, and the WEHI computing cluster.

The AI tool Claude 3.5 Sonnet (Anthropic, 2024) was used for proofreading the manuscript and checking for consistency in spelling, grammar, and clarity of the text. It was not used for the generation of ideas, content, figures, or data.

## Author contributions

Conceptualization: KAB, AG

Methodology: KAB, JIM, HV, TAD, AL, TAR, LLLW, AG

Investigation: KAB, JIM, HV, TAD, AL, TAR, LLLW, AG

Funding acquisition: DMT, AG

Supervision: LFD, DMT, AG

Writing – original draft: KAB, AG

Writing – review & editing: All authors

