## Supplemental figures for "Structural basis for Porcupine inhibition"

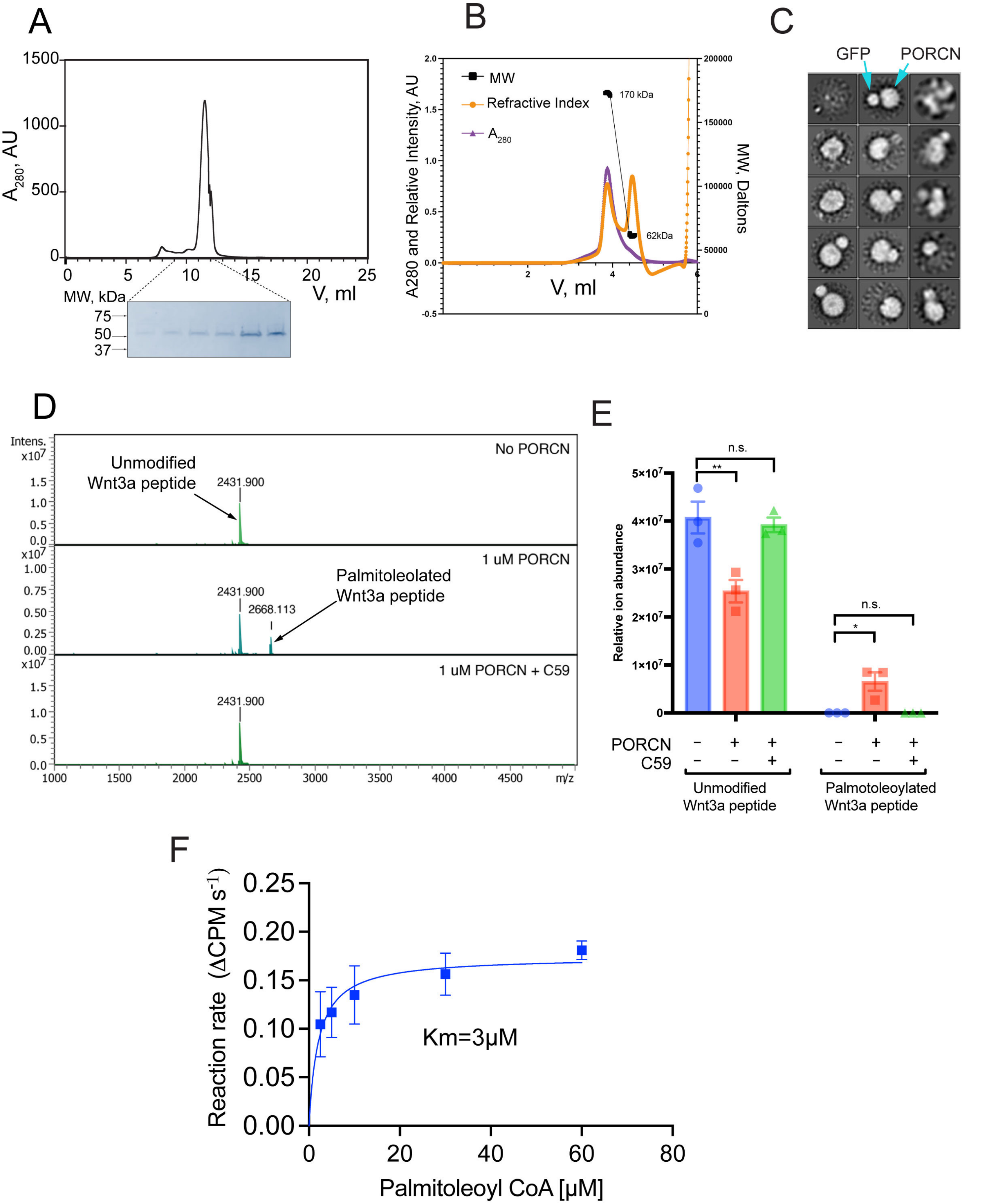


**Supplementary Figure 1. Purification and functional characterisation of PORCN. A)** Size exclusion chromatography profile of purified PORCN with SDS-PAGE analysis of peak fractions (inset). **B)** SEC-MALS analysis of PORCN showing a monodisperse peak at 170 kDa. **C)** Representative negative stain EM images of purified PORCN-GFP. **D)** LC-MS analysis of PORCN-catalysed palmitoylation of Wnt3a peptide in the absence (middle) or presence (bottom) of C59 inhibitor, compared to no PORCN control (top). **E)** Quantification of LC-MS data (n=3, mean ± SEM, One-way ANOVA, *P < 0.05, **P < 0.01, n.s. not significant). **F)** Kinetics of PORCN-dependent Wnt3a peptide acylation. Data fitted to the Michaelis-Menten equation ($K_m$ = 3 μM).

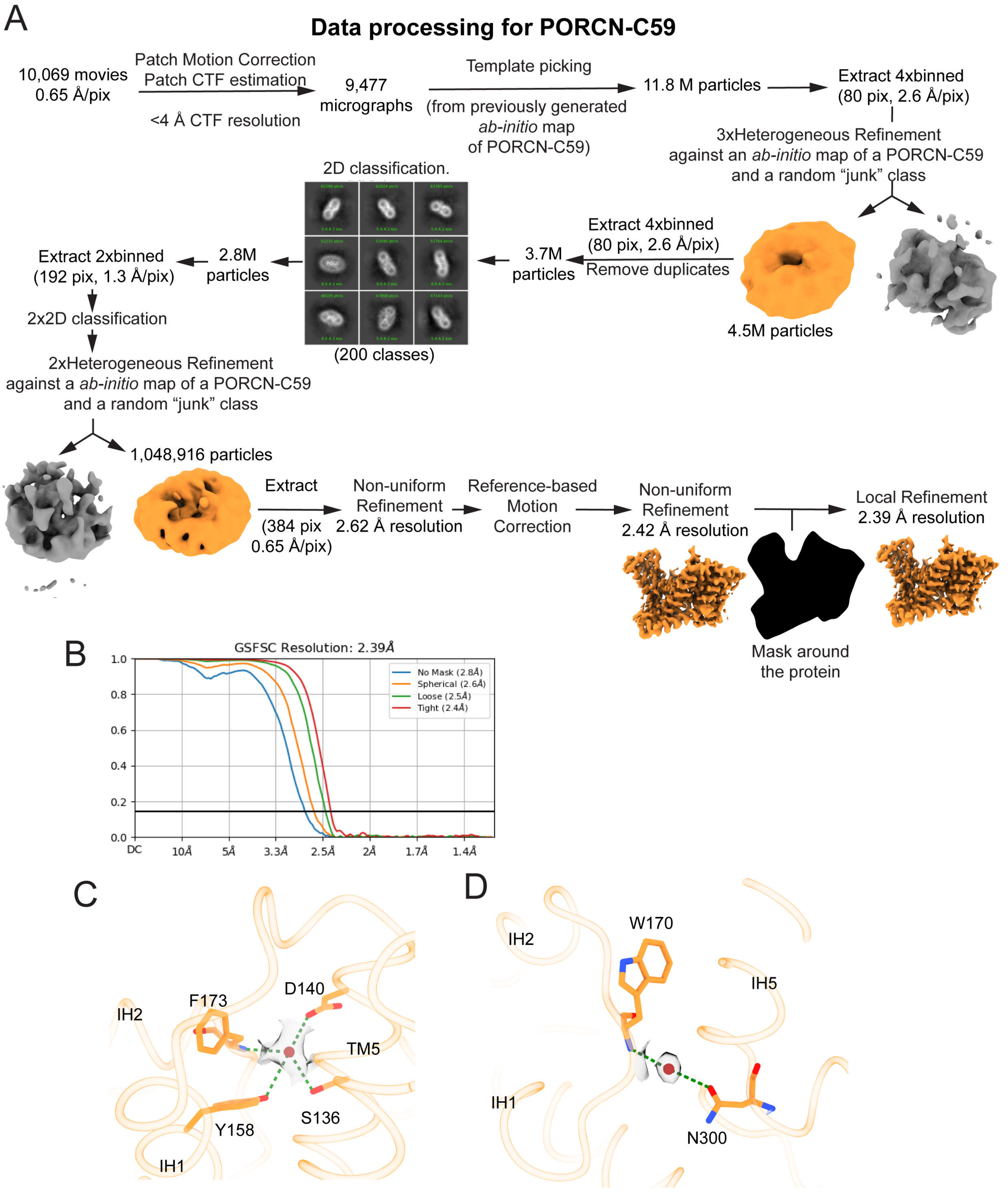


**Supplementary Figure 2. Cryo-EM data processing of PORCN-C59 complex. A)** Data processing workflow for PORCN-C59 complex. **B)** Gold-standard FSC curves for the final 3D reconstruction with different masking strategies. **C-D)** Cryo-EM density (transparent grey surface, map contour level 0.02) for the structural water molecules (red spheres) in PORCN-C59 structure. Coordinating residues are shown as sticks. H-bonds are shown as green dashed lines.

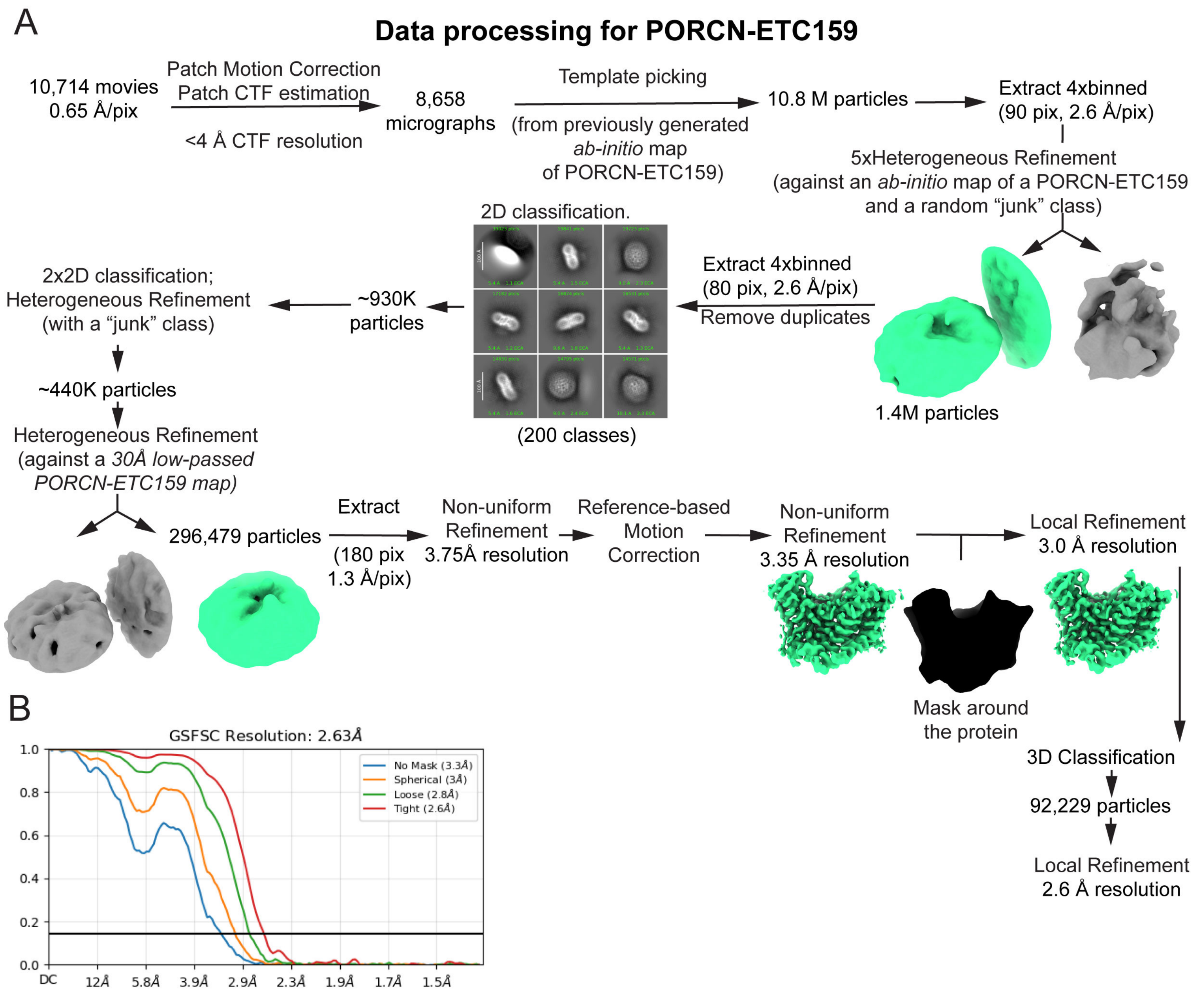


**Supplementary Figure 3. Cryo-EM data processing of PORCN-ETC159 complex. A)** Data processing workflow for PORCN-ETC159 complex. **B)** Gold-standard FSC curves for the final 3D reconstruction with different masking strategies.

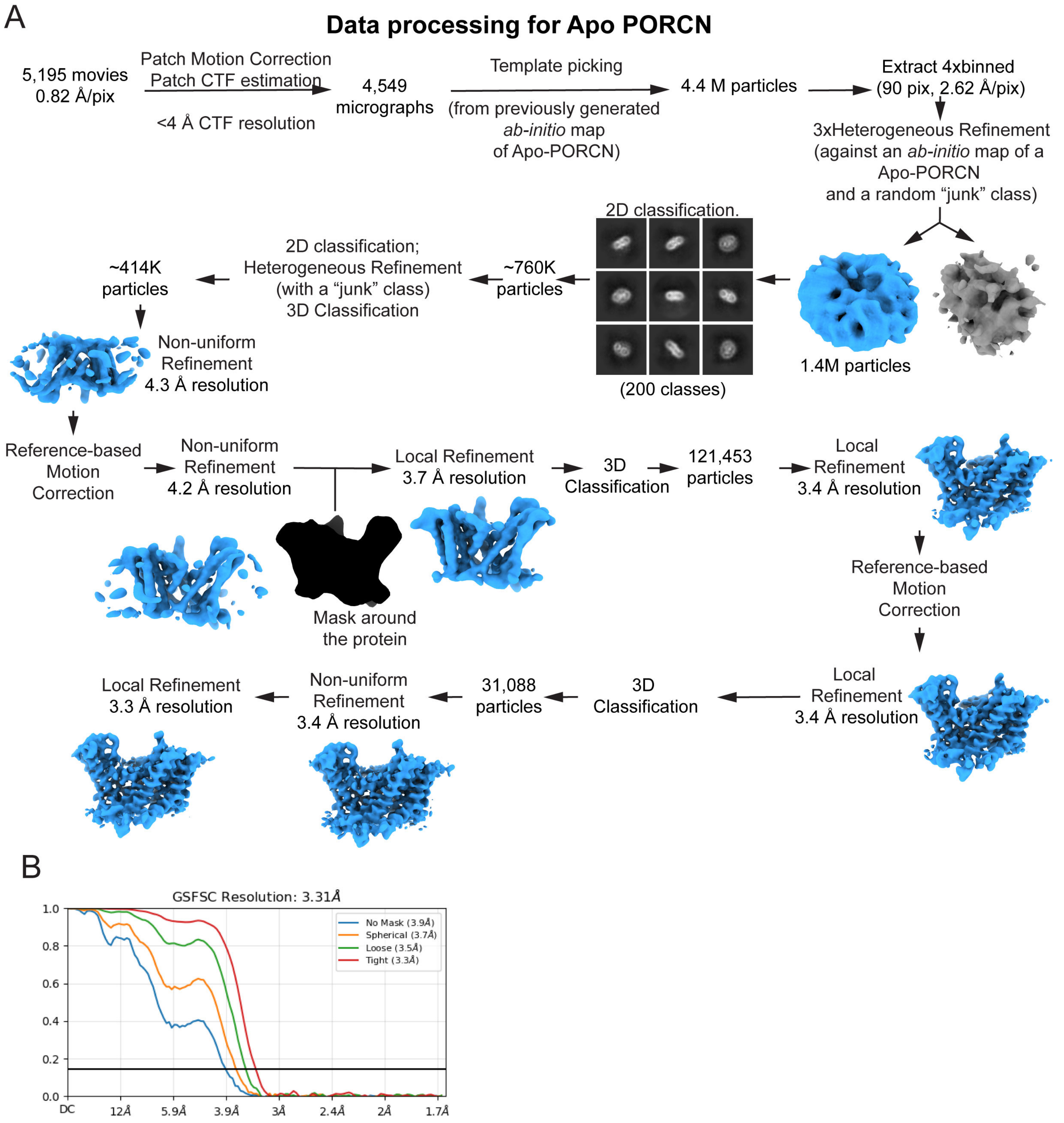


**Supplementary Figure 4. Cryo-EM data processing of apo-PORCN. A)** Data processing workflow for apo-PORCN. **B)** Gold-standard FSC curves for the final 3D reconstruction with different masking strategies.

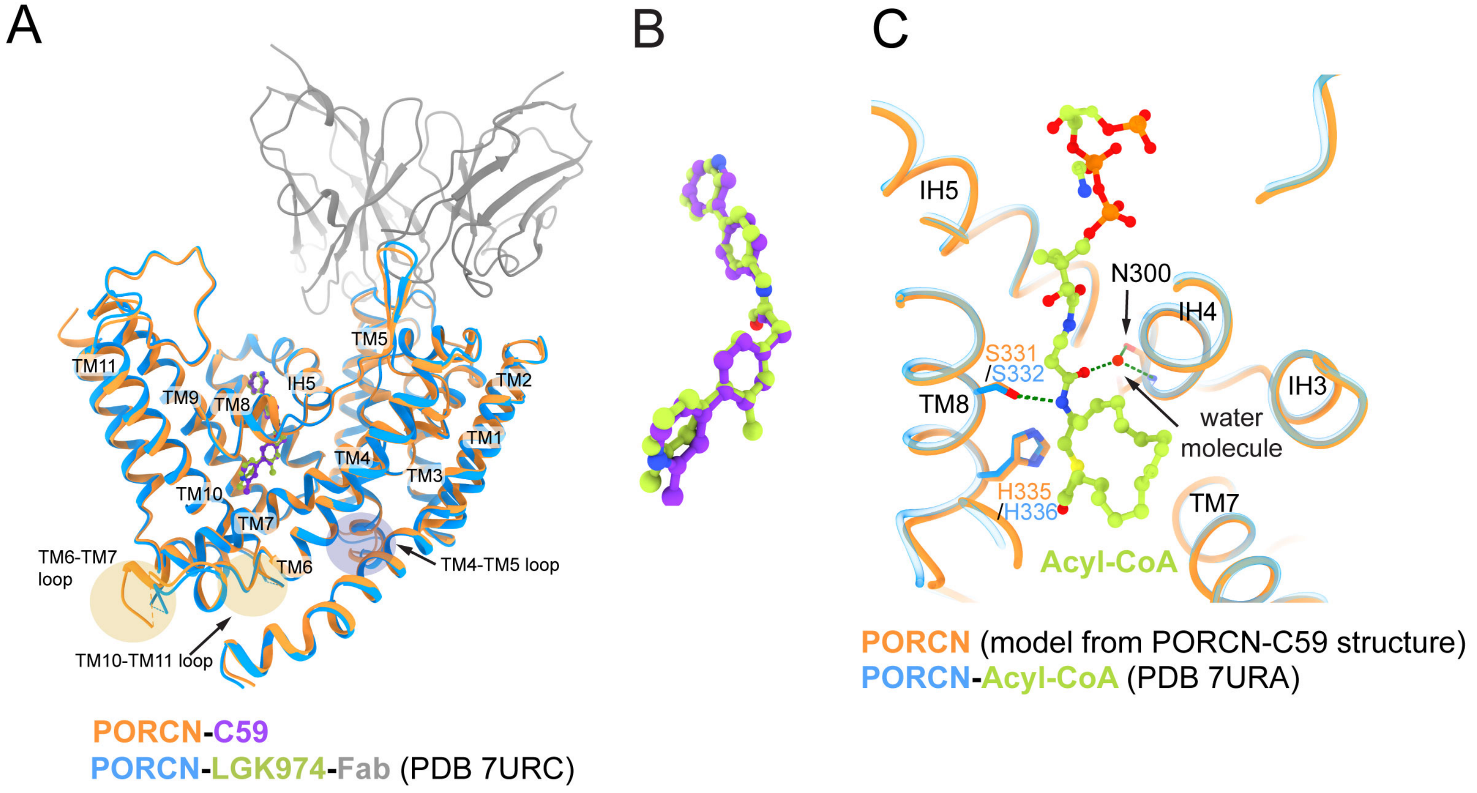


**Supplementary Figure 5. Structural comparison with published PORCN structures. A)** Superposition of PORCN-C59 (orange-purple) and PORCN-LGK974-Fab (PDB 7URC, blue-light green-grey) structures. **B)** Close-up view of superimposed C59 and LGK974. **C)** Overlay of the PORCN-acyl-CoA (PDB 7URA, blue-light green) with the PORCN-C59 structures (C59 omitted for clarity) showing that the water molecule (red sphere) observed in our PORCN-inhibitor structures is important for positioning the acyl chain for catalysis.

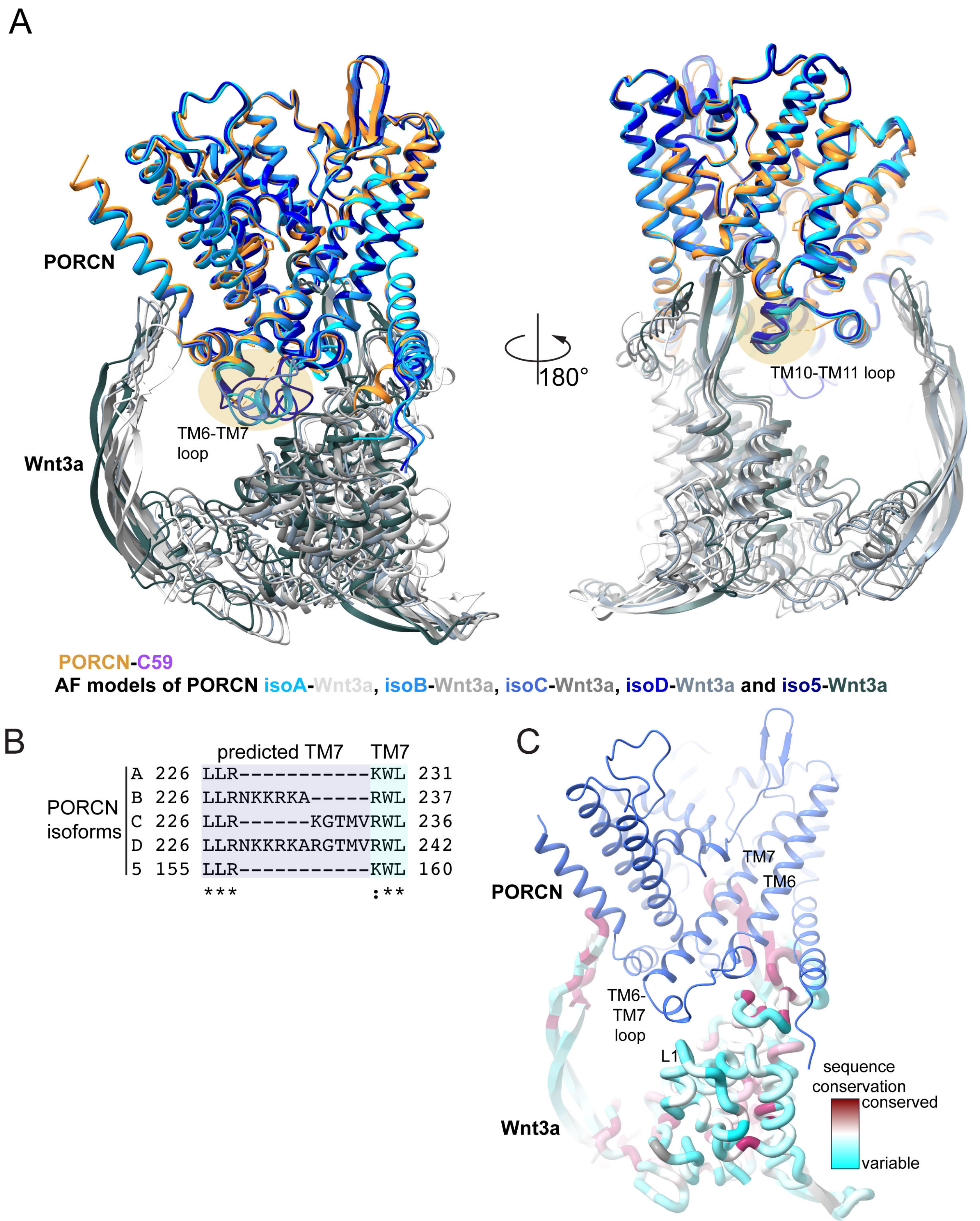


**Supplementary Figure 6. Models of 5 PORCN isoforms and their predicted interactions with Wnts. A)** AlphaFold3-generated models of five PORCN isoforms in complex with Wnt3a superposed with the PORCN-C59 structure. The ER lumen-facing loops (TM6-TM7 and TM10-TM11) that are unresolved in our structures are predicted to interact with Wnt. **B)** Sequence alignment of five human PORCN isoforms showing the alternatively spliced region at TM7. **C)** Model of PORCN-Wnt3a complex coloured by sequence conservation among human Wnts, highlighting the variable L1 loop region in Wnt that may contribute to PORCN isoform selectivity.

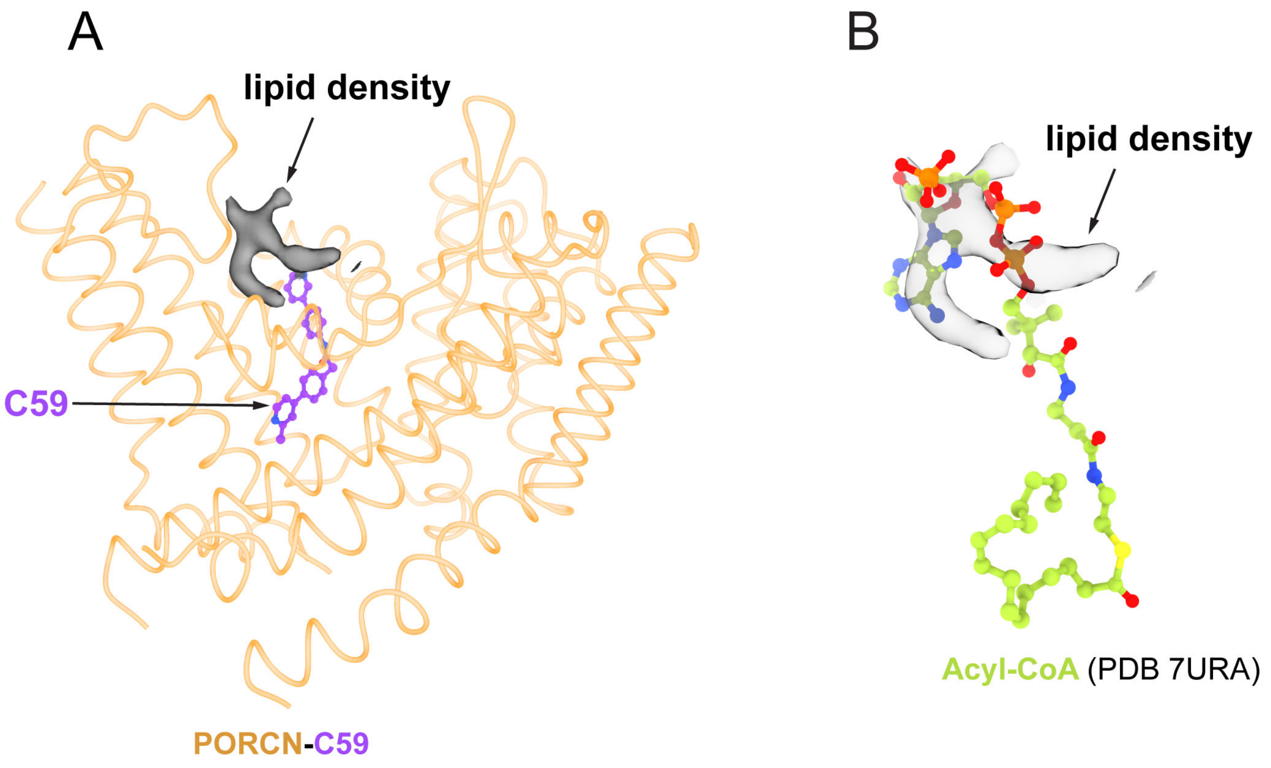


**Supplementary Figure 7. Lipid-like density at the entrance to the PORCN inhibitor binding site. A)** Unassigned lipid-like density (transparent grey surface, map contour level 0.048) observed at the cytosolic entrance of the PORCN-C59 binding pocket. **B)** Superposition with acyl-CoA from PDB 7URA (light green) showing the lipid-like density overlapping with the phospho-adenosine headgroup of acyl-CoA.

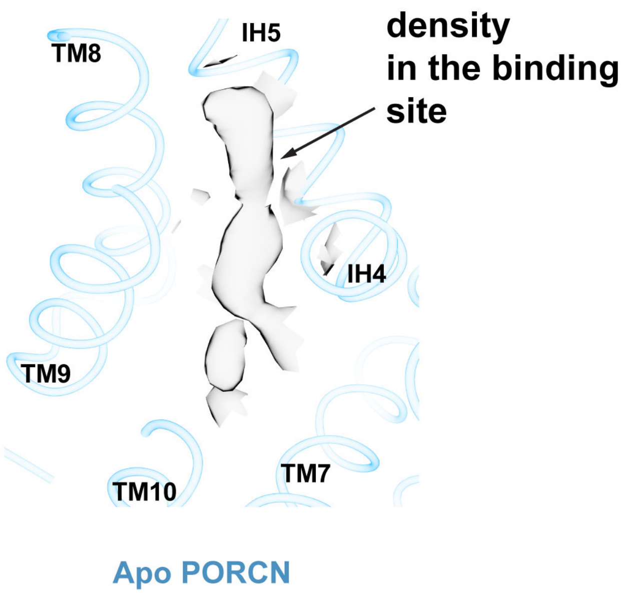


**Supplementary Figure 8. Electron density in the binding site of apo-PORCN.** Weak electron density (transparent grey surface, map contour level 0.034) observed in the binding site of apo-PORCN, which could represent water molecules or low-occupancy endogenous ligands.

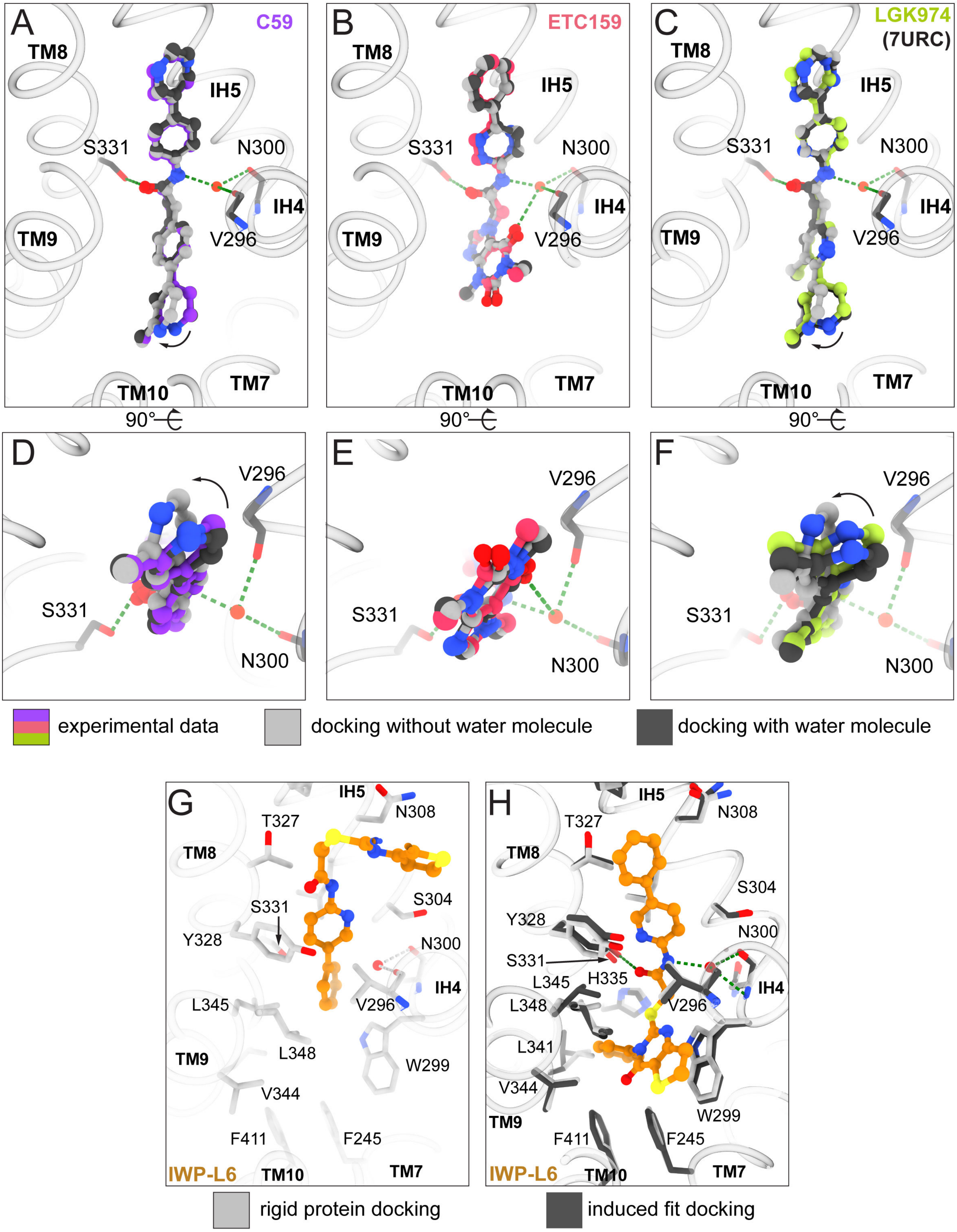


**Supplementary Figure 9. Docking validation and analysis of IWP-L6 binding. A-F)**, Comparison of experimentally determined (purple, pink or light green) and docked (grey or dark grey) poses of C59, ETC159, and LGK974, showing improved accuracy when the water molecule is included in the docking simulations. **G)** Initial rigid protein docking of IWP-L6 (orange) showing binding at the entrance of the binding site. **H)** Induced fit docking of IWP-L6 showing the expected binding mode after allowing protein side-chain flexibility.
