## Supplemental tables for "Structural basis for Porcupine inhibition"

**Table S1. Cryo-EM data collection, refinement and validation statistics**

| | PORCN-C59 (EMDB 70660) (PDB 9OO6) | PORCN-ETC159 (EMDB 70661) (PDB 9OO7) | Apo PORCN (EMDB 70662) (PDB 9OO8) |
|---|---|---|---|
| **Data collection and processing** | | | |
| Magnification | 130,000x | 130,000x | 105,000x |
| Voltage (kV) | 300 | 300 | 300 |
| Electron exposure (e–/Å$^2$) | 60 | 70 | 60 |
| Defocus range (μm) | 0.5-1.5 | 0.5-1.5 | 0.5-1.5 |
| Pixel size (Å) | 0.65 | 0.65 | 0.82 |
| Symmetry imposed | C1 | C1 | C1 |
| Movies collected (no.) | 10,069 | 10,714 | 5,195 |
| Initial particle images (no.) | 11.8M | 10.8M | 3.8M |
| Final particle images (no.) | 1,047,762 | 92,229 | 31,088 |
| Map resolution (Å) 0.143 FSC threshold | 2.39 | 2.61 | 3.32 |
| Map resolution range (Å) | 2.0-2.6 | 2.3-3.2 | 2.9-4.0 |
| **Refinement** | | | |
| Initial model used (PDB code) | AlphaFold2 PORCN model | PDB 9OO6 | PDB 9OO6 |
| Model resolution (Å) 0.143 FSC threshold | 2.33 | 2.51 | 3.27 |
| Model resolution range (Å) | 2.1-2.8 | 2.2-3.7 | 3.0-4.4 |
| Map sharpening *B* factor (Å$^2$) | -60 | -30 | -50 |
| Model composition | | | |
| Non-hydrogen atoms | 3610 | 3484 | 3378 |
| Protein residues | 438 | 431 | 423 |
| Ligands | 3 | 2 | 1 |
| *B* factors (Å$^2$) | | | |
| Protein | 90 (58-172) | 30 (5-107) | 44 (11.7-120) |
| Ligand | 104 | 58 | 134 |
| R.m.s. deviations | | | |
| Bond lengths (Å) | 0.004 | 0.004 | 0.004 |
| Bond angles (°) | 0.733 | 0.766 | 0.722 |
| Validation | | | |
| MolProbity score | 1.25 | 1.21 | 1.37 |
| Clashscore | 4.85 | 4.31 | 5.3 |
| Poor rotamers (%) | 0.27 | 0 | 0 |
| Ramachandran plot | | | |
| Favored (%) | 98.4 | 98.6 | 97.6 |
| Allowed (%) | 1.6 | 1.4 | 2.4 |
| Disallowed (%) | 0 | 0 | 0 |

**Table S2. Docking scores for PORCN inhibitors**

| Ligand | Docking scores (rigid protein) | |
|---|---|---|
| | No water | With water |
| C59 | -10.5 | -11.1 |
| ETC159 | -9.9 | -9.5 |
| LGK974 | -7.6 | -6.1 |
| GNF6321 | -8.2 | -8.9 |
| RXC004 | -9.3 | -11.0 |
| IWP-O1 | -8.9 | -10.1 |
| IWP-L6* | -8.9 | -7.1 |

*The IWP-L6 did not bind in the expected pose using the rigid docking procedure
